# Plant Acupuncture: a low-cost and open-source device for local mechanical stimulation

**DOI:** 10.64898/2026.09.10.750389

**Authors:** Anna Daamen, Annalisa Bellandi, Maarten Besten, Jasper Lamers, Sjoerd Woudenberg, Clint Heijstek, Jan Willem Borst, Cecilia Borassi, Joris Sprakel

## Abstract

**Background:** Mechanical signals are important regulators of cellular responses in plants. They guide plant development and can activate defense and repair mechanisms. Yet, the molecular mechanisms by which plants perceive, transduce, and interpret mechanical signals are still poorly understood. This is in part due to the lack of methods to apply local, precise and non-damaging mechanical forces to plant cells. Micro-indentation is highly suitable for this purpose, yet available instrumentation is often expensive and difficult to combine with high-resolution microscopy.

**Results:** We designed an open-source and affordable modular indentation device, consisting of 3D printed elements, 3 commercially available piezo motors, and a variety of indentation needles. Due to its modularity, the setup can be readily adapted to meet experimental requirements and works on all microscopes with bespoke adaptors. We show that the setup can be used to explore both rapid and slower touch responses, exemplified by visualizing calcium waves and actin patches induced by touch. We also show that the setup is compatible with various plant species and tissues and can be combined with high-resolution functional imaging.

**Conclusions:** The simple and flexible design of the indentation device presented in this paper ensures that any lab with a 3D printer can build their own setup at low cost and with minimal time investment. The system has a wide range of applications for live plant tissue, making indentation experiments and thereby plant mechanobiology studies, more accessible.

## Background

All living organisms are continuously exposed to mechanical forces. From the touch we experience as human beings, to the adhesive forces bacteria sense when growing on a surface (3-5), to the wind blowing against a tree (6, 7). The capacity to sense mechanical forces is a universal feature across the entire tree of life. However, the fact that these mechanical forces can act as signals triggering biochemical and genetic responses in cells has only been appreciated in the past few decades (8-12). Especially in plants, the molecular mechanisms by which cells or tissues sense forces and mechanical properties (mechanosensing) and transduce these into biological responses (mechanotransduction) are poorly understood.

For plants, mechanical signals can originate from the environment, such as wind (6, 7), the soil they grow through (13), and the enormous local forces generated during pathogen invasion (14-18), or from within, such as turgor pressure (19, 20), differential growth of tissues and differential stress patterns in tissues due to inhomogeneous cellular morphology (12, 20-23). All these mechanical signals are important regulators of cellular and genetic processes in plants, both in reproduction (24, 25), during defense responses to pathogens or wounding (16-18, 26), and in morphogenetic processes in tissue development (12, 20-23, 27-30).

To study mechanobiological regulation, various options exist in order to apply well-defined mechanical stimuli to plants. Osmotic treatments are widely used to manipulate cellular turgor pressure by changing extracellular osmolarity (12, 31-33). The outward-directed turgor pressure is mechanically balanced by the inward-oriented elastic restoring force of the turgor-stretched cell wall. This places plant cells in a state of mechanostasis, governed by a balance between compressive and tensional forces, a so-called tensegrity balance. Changing extracellular osmolarity disturbs the tensegrity balance and, in turn, alters the tension on cell walls and membranes. Osmotic stress acts globally across the entire tissue or organism submerged in the medium; hence, it is not suitable for studying very local responses. Another method of applying mechanical stress to an entire tissue is to compress or stretch it. Compression is usually applied using coverslips or relatively large compression probes (29, 33-35). Stretching assays have been implemented in several ways, including fixing one side of the tissue and attaching the other to a weight, clamping tissues between independently movable plates, or mounting the tissue onto an elastomeric substrate that is subsequently stretched (36-38). While these strategies allow quantitative application of a strain to an entire tissue, this can still result in an inhomogeneous mechanical field in the sample, as the distribution of mechanical stresses is highly dependent on the local geometry and mechanics of the tissue. As such, these methods are not suitable for local mechanical stimulation to explore sub-cellular and cell polarity responses.

If a local mechanical signal is needed, the most common technique is to deflate cells or tissue sections, either using laser ablation (18, 22), needle-based ablation (36), or incisions (29). These techniques cause local damage to one or a few cells, thereby releasing the turgor pressure of those cells. Deflating a single cell causes its neighboring cells to lose counterpressure at one face. As a result, the tissue-wide tensegrity balance is disturbed, leading to a redistribution of the stress pattern. Though effective and highly local, these techniques cause substantial damage. Damage leads to the release of damage-associated molecular patterns (DAMPs), which can be cell wall fragments, peptides or cell-internal compounds such as ATP or DNA (39, 40). These DAMPs can trigger damage response pathways that are not necessarily caused by mechanical stresses (41). This makes it difficult to distinguish mechanical from biochemical damage signaling. An alternative for local, non-damaging application of forces is to indent plant cells from the outside using a needle (35, 42, 43). Indentation of plant cells results in highly localized deformation of the cell wall and allows the controlled application of a force on the cell. This makes indentation the only technique that achieves targeted, localized deformation with high accuracy, without causing tissue damage. In turn, this makes it possible to separate DAMP signaling from true mechanical signaling. However, indentation has remained less widely adopted than other mechanical stimulation techniques, mostly due to practical limitations of available setups. Users are often limited to the setup as it is, without the ability to alter it as needed, for example, to fit a different microscope or to extend its capabilities. Most of all, indentation setups can be expensive and are often not designed for use with high-resolution fluorescence or functional microscopy.

To facilitate broader application, here we report the development of a modular, open-source indentation device optimized for live plant tissues. The system is designed to be (i) cost-effective and easy to assemble using 3D-printed components combined with a minimal set of commercially available parts; (ii) easily adaptable to different microscopes and to experimental requirements for high-resolution imaging.

## Results

### Design of the device

The aim of this study is to develop a device that enables local mechanical manipulation of plant cells, specifically soft, non-puncturing indentations, while simultaneously performing high-resolution live imaging. In addition, we want the design to be compatible with a variety of microscopes via interchangeable mounts and flexible to meet specific needs, such as puncturing rather than indentation. Moreover, in line with the open-science principle, we want the device to be affordable and easy to recreate in any lab without requiring advanced technical expertise. To meet these criteria, we designed a device consisting of three small, commercially available, affordable piezo motors mounted in a 3D-printed assembly.

The motor block for moving the indentation needle contains a dual piezo motor for movements in the x-y plane, parallel to the imaging plane upon mounting the device onto a microscope. Using a right-angle bracket, another piezo motor is mounted vertically for movements along the z-axis, perpendicular to the imaging plane (Figure 1a-d, Figure S1a-c). Each piezo motor is connected to a controller, which is linked to a PC for control via the manufacturer-supplied software (Supplementary Figure 1a). Coarse positioning is achieved using the joystick on the controller, while fine positioning with micrometer precision is controlled via the software on the computer.

**Figure 1.**
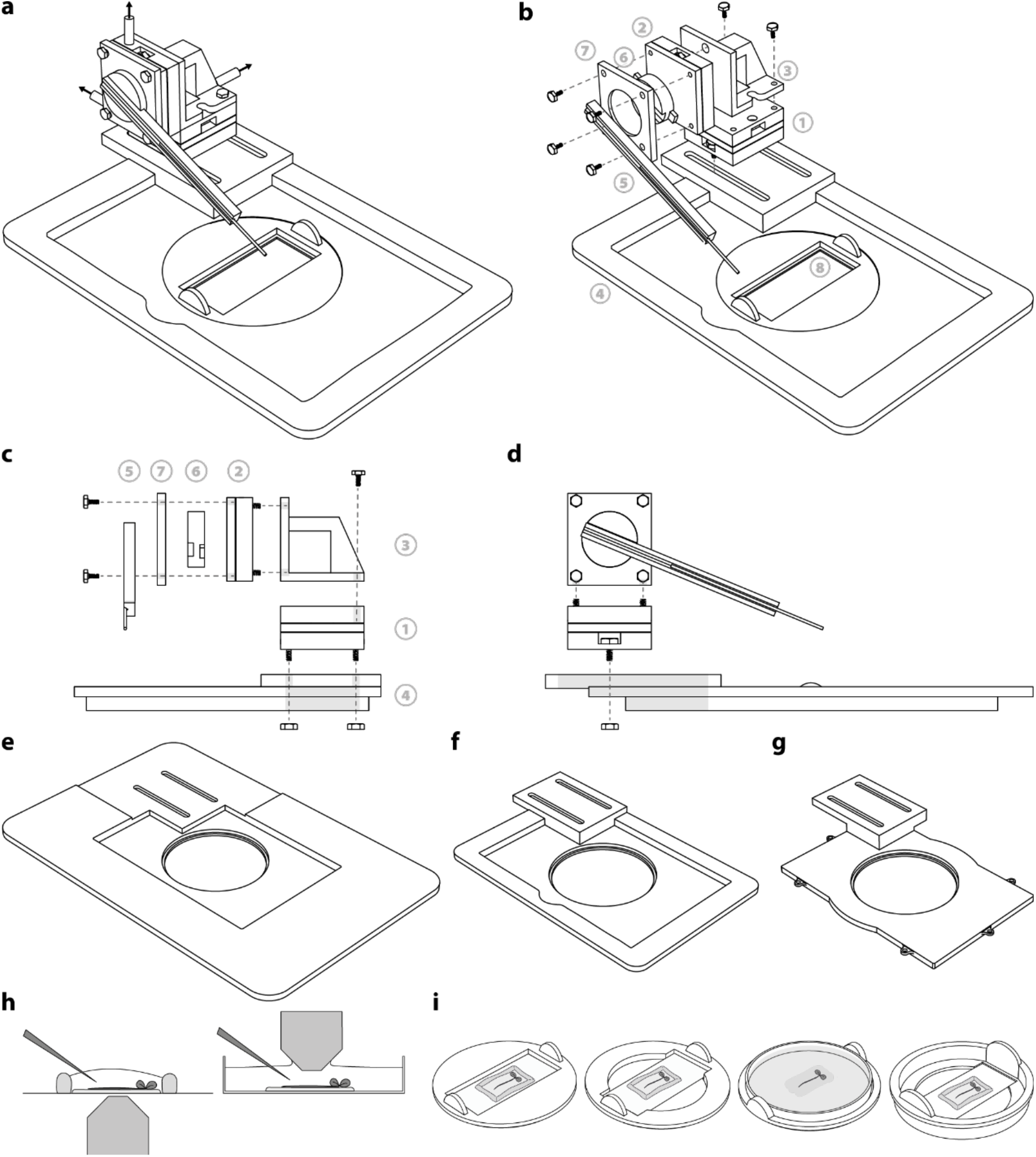
Design of the indentation device. **(a-d)** The motor block for moving the needle consists of a horizontal dual motor (1) and a single motor (2), mounted vertically to a right-angle bracket (3). This block is mounted onto a 3D printed base (4) that is designed to fit into the microscope. Each piezo motor has an outlet that can be connected to the controller to move each motor individually (outlets indicated by the arrows in **a**). The needle is glued onto a thin 3D printed rod with a slit for the needle (5). This rod clicks into a round rotating needle holder (6) that is mounted onto the vertical piezo motor with a 3D printed mount (7). Clicking enables easy exchange of needles. The sample is placed into the base on a sample holder (8). **(b-d)** Exploded views of the setup. **(e-g)** Bases designed for Nikon **(e)**, Leica SP8 DIVE **(f)**, and Leica SP5 and SP8 **(g)** microscopes. **(h)** Cross section of mounted samples. The seedling is glued onto a cover glass or petri dish using KwikSil low toxicity glue and submerged in medium. Due to the transparency of the glue, the sample can be imaged with inverted microscopes (left), or an upright microscope that has a water immersion objective (right). **(i)** Samples are placed into a sample holder. From left to right: a stable holder supporting the whole cover glass, a holder that leaves more space for large objectives in inverted microscopes, a holder that fits a 6 cm petri dish for up-right microscopes with a dipping objective, a holder where the cover glass is lowered to move it closer to the objectives. The sample holder is placed into the base of the device and can be rotated in any desired position.

The indentation setup controls a thin needle that contacts the sample. The system is compatible with a wide range of needles (discussed below in more detail) and can be mounted to thin 3D-printed rods with a slit to securely hold the needle (Figure1a-d). These rods are clicked into a rotating needle holder mounted onto the vertical piezo motor with a 3D-printed mount (Figure 1a-d, Figure S1b). This click-in system enables rapid and easy exchange of needles between or during experiments. Moreover, the whole motor block with needle can be mounted onto different stage insert bases that are tailored to specific microscopes, here demonstrated for Nikon microscopes (Figure 1e, Figure S1d), Leica SP8 DIVE (Figure 1f, Figure S1e), and Leica SP5 and SP8 systems (Figure 1g, Figure S1f). Full design files for 3D printing these parts are available with this paper as a supplement and allow anyone to build this device themselves. In total, the costs of building the setup are approximately €3600, including all electronic elements and the material for 3D printing. Needles cost between €10 for acupuncture needles to €75 for tungsten needles.

To indent exterior cells of a plant, the sample must be fixed in place. If this is not the case, the sample will be pushed away by the needle rather than being indented. To this end, we glue plant tissues onto a thin layer of Kwik-Sil silicone glue that is designed for biological samples (Figure 1h, Figure S1g). A reservoir for medium can be created around the tissue with vacuum grease. The sample is adhered onto a cover glass, which is then placed into a 3D-printed sample holder (Figure 1h, Figure S1h-k). This sample holder can rotate when placed in the base, allowing the sample to be oriented in any preferred way with respect to the needle and imaging set-up. Four different sample holders have been designed: a stable holder that supports the entire cover glass (Figure 1i, Figure S1h), a holder that allows for more space for large immersion objectives (Figure 1i, Figure S1i), a holder that lowers the sample more towards the objective in case the microscope has a limited working distance (Figure 1i, Figure S1j) and finally a holder that fits a small petri dish for water dipping objectives (Figure 1i, Figure S1k). The variety in sample holders and bases ensures that the setup fits on many microscopes. As the glue is transparent, it is suitable for both inverted microscopes and upright microscopes using dipping objectives.

### Needle geometry

Since the needles are fixed onto 3D-printed holders, the setup is compatible with virtually any type of needle. To assess the performance of different needle types, we tested five different needles: commercially available tungsten needles with a 1 µm and 5 µm tip diameter, commercially available steel acupuncture needles with a tip diameter of 8 µm, and custom-made glass needles with tip diameters of 12 and 18 µm (Figure 2a). We performed indentations on epidermal cells in the maturation zone of *Arabidopsis thaliana* seedling roots. To visualize the cell wall during indentation, the samples were stained with propidium iodide (PI).

**Figure 2.**
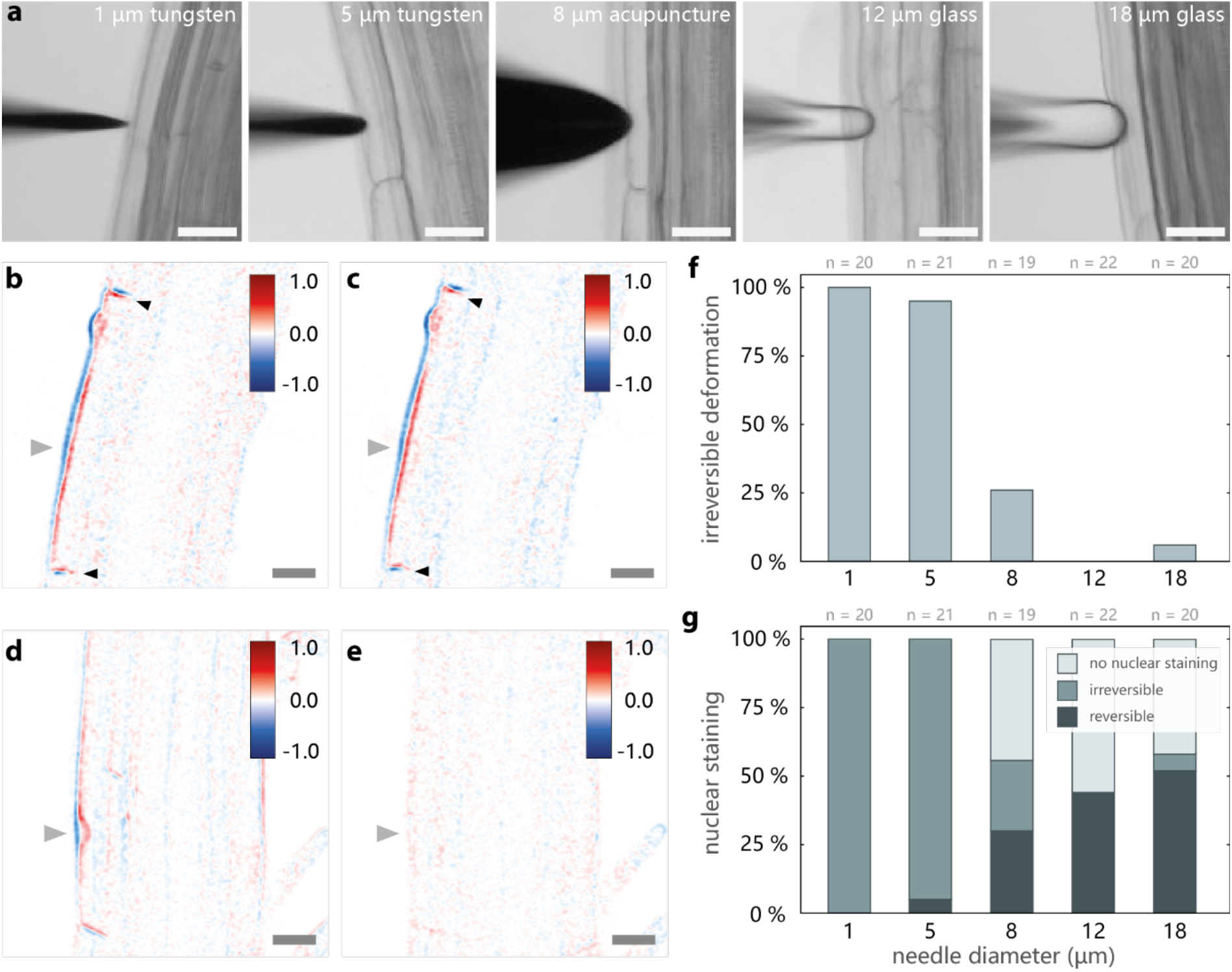
Effect of needle geometry on the indented sample. **(a)** Needles used in this study. From left to right: tungsten needle with 1 µm tip diameter, tungsten needle with 5 µm tip diameter, acupuncture needle with 8 µm tip diameter, glass needle with 12 µm tip diameter, glass needle with 18 µm tip diameter. **(b-c)** Difference maps of a cell that is indented with a 1 µm tungsten needle. Grey arrow indicates the position of the needle. The color bar shows normalized intensity, where negative values indicate signal that disappeared/decreased and positive values indicate signal that appeared/increased. Difference maps show deformation during indentation **(b)** and after indentation **(c)**. Black arrows highlight cell walls that typically move inward when the cell loses turgor pressure. **(d-e)** Difference maps of a cell that is indented with an 18 µm glass needle. Difference maps show deformation during indentation **(b)** and after indentation **(c). (f)** Percentage of irreversible deformations during indentations with the 5 different needles. **(g)** Quantification of nuclear staining caused by PI internalization in the indentations in **f**. Scale bars are 25 µm.

Cell deformations created by the needle were visualized by differential microscopy (DM). This is a data analysis approach in which two images, first registered to align them as closely as possible, are subtracted from each other. Areas in which cell walls have moved appear as either positive or negative values, while areas that remain unchanged disappear from the difference map as their difference is zero. Two DM comparisons were made: 1) before indentation versus during cell indentation, showing the extent of deformations during contact, 2) before indentation versus after retracting the needle, revealing any permanent changes in the cell shapes resulting from the indentation, i.e., verifying reversibility. For example, with a 1 µm tungsten needle, inward movement of the cell wall during contact (Figure 2b) remained after retracting the needle (Figure 2c). This permanent deformation and the inward movement of the cell walls indicate a loss of turgor pressure due to the indentation and thus reveals that this indentation was irreversible due to cell puncture. By contrast, a larger, rounded needle, such as the glass needle with an 18 µm tip diameter, shows a highly local deformation of the cell wall during contact (Figure 2d), yet this deformation is fully reversible as no permanent deformation is visible after retracting the needle (Figure 2e).

Quantitative analysis of DM data shows that the tungsten needles with a small tip diameter almost always cause irreversible deformations of the cells in *A. thaliana* roots (100% for a 1 µm tungsten needle and 95% for a 5 µm tungsten needle; Figure 2f). The 8 µm acupuncture needle caused irreversible deformations in only 26% of the indentations, while most deformations were reversible. Glass needles with a larger tip diameter predominantly produced soft, reversible deformations of the cells. Only a single indentation with the 18 µm glass needle resulted in mild irreversible deformation, likely due to a greater indentation depth compared to other indentations. We used PI as an indicator of cell damage, as damaged cells become permeable to PI, allowing it to enter the cell and stain DNA in the nucleus. For indentations annotated as irreversible according to our DM analysis, this was indeed the case. The nucleus is stained in 100% of the indentations with both 1 µm and 5 µm tungsten needles (Figure 2g). For the 8 µm acupuncture needle, the nucleus was always stained when the deformation was irreversible. Yet, in some cases, the nucleus was also stained when the cell deformation was reversible. Nuclear staining was observed in 30%, 44%, and 52% of the reversible indentations with 8 µm acupuncture, 12 µm glass, and 18 µm glass needles, respectively. As the frequency of nuclear staining increased with tip diameter, we hypothesize that membrane stretching during indentation can transiently permeabilize the membrane, allowing PI to enter the cell.

We consistently observe a release of turgor pressure and irreversible deformation of neighboring cells, bulging toward the indented cell, with the 1 µm tungsten needle. This is like what is described for laser ablated cells (12, 44), and as such we interpret this as cells being punctured and deflated due to the indentation. As such, for experiments in the next sections, this specific needle is used to puncture cells. In contrast, reversible indentations are reproducible with the glass needles with a larger diameter. Since we observe transient permeabilization of the membrane when this needle is used, this implies that even reversible deformations of cells can lead to temporary membrane permeabilization, which is a weaker and temporary form of mechanical damage as compared to that experienced during cell puncture and lysis.

### High-resolution imaging of sub-cellular touch responses

We developed the device to enable high-resolution live imaging of cellular responses to local touch. A key rapid cellular response to mechanical stimulation is the generation of a traveling calcium wave that spreads from the point of contact (45-47). We indented plants that express the fluorescent calcium marker *p35S::R-GECO1*.*2* to verify if rapid responses can be monitored with our device. Because calcium dynamics occur on a second timescale, capturing these dynamics requires imaging at high temporal resolution while the indentation device is in operation, i.e., while the needle moves toward and away from the sample. Since we use piezo motors that move smoothly, this works well in our setup. As anticipated, a calcium signal emerges within 5 seconds after touch (Figure 3a). Interestingly, we find that the signal amplitude and its propagation depended on the needle geometry. With a 13 µm glass needle, which results in reversible indentations, a calcium signal emerges only in the cell that is indented. The signal disappears within 20 seconds after indentation (Figure 3a, Additional File 2). By contrast, puncturing cells with a 1µm tungsten needle triggers a higher-amplitude calcium wave that spreads to neighboring cells within 20 seconds and then fades (Figure 3a, Additional File 3). While both indentations have a mechanical component, the cellular responses are distinct when comparing an indentation that causes damage to one that is purely mechanical. These findings demonstrate that the device enables real-time visualization of rapid and dynamic cellular touch processes. Moreover, the ability to change needles and thereby precisely control the mechanical stimulus allows discriminating between mechanical and damage signaling within a single approach.

**Figure 3.**
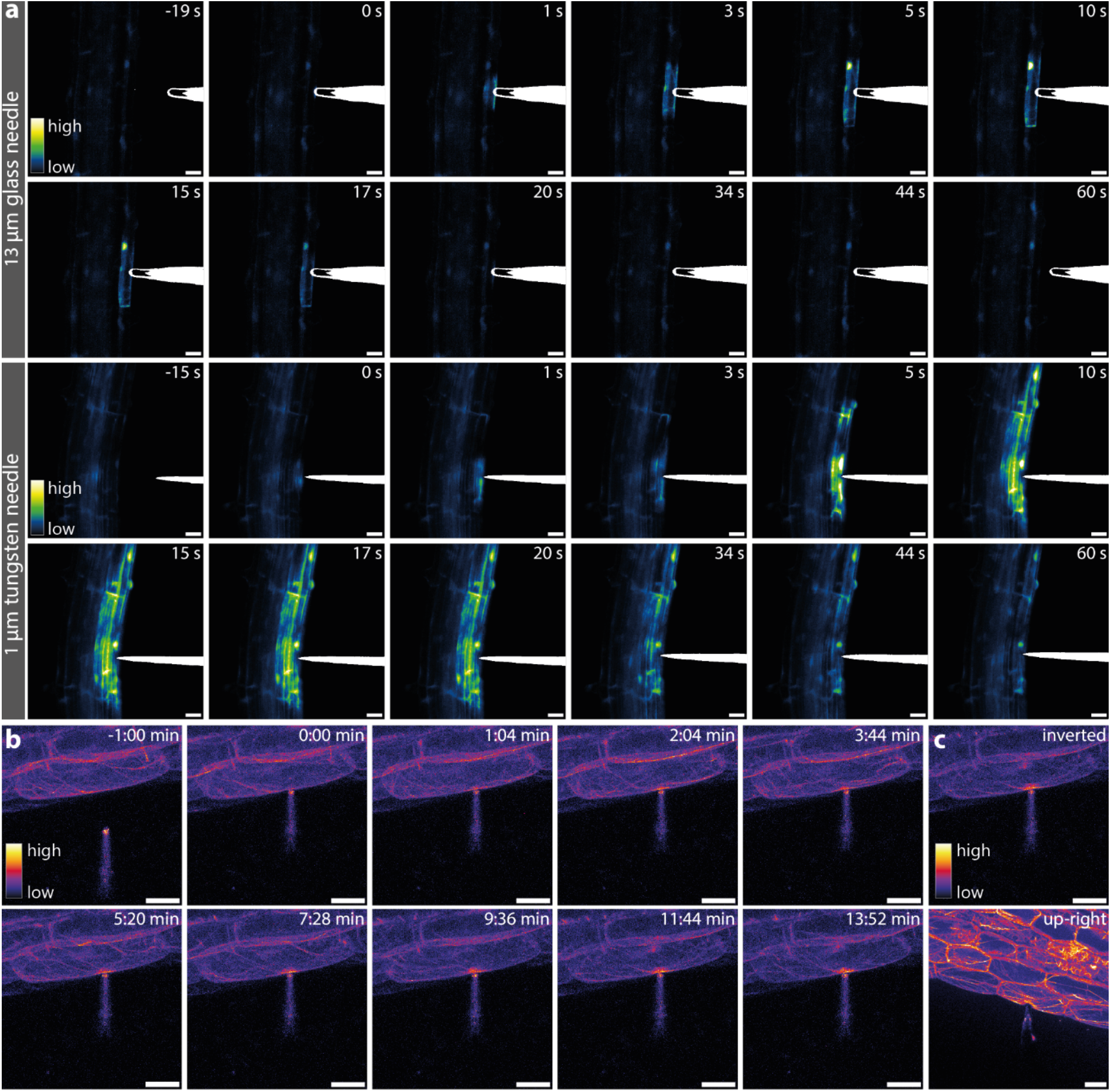
Indentations on A. thaliana roots enable studies of dynamic processes. **(a)** Timeseries of reversible indentation with a 13 µm glass needle and puncturing of a cell with a 1 µm tungsten needle on cells in the maturation zone of *A. thaliana p35S::R-GECO1*.*2* roots. Reversible indentation confines the response to the indented cell only, while cell damage enables propagation of the calcium signal to neighboring cells. Brightness of the images with the glass needle has been increased with a factor 2.7. This was not necessary for the images with the tungsten needle. **(b)** Maximum projections of indentation of a hypocotyl cell of *A. thaliana pUBQ10::ABD2-mCherry* seedling with a 5 µm tungsten needle. Indentation of the cell induced actin accumulation to the indentation side within 2 minutes, which accumulates to form a patch over 14 minutes. **(c)** Maximum projections of indentations of a hypocotyl cell of *A. thaliana pUBQ10::ABD2*-mCherry with a 5 µm tungsten needle. Comparison of an image acquired with an inverted microscope with a water immersion objective (40X NA 1.10) (top) to one that is acquired with an upright microscope equipped with a water dipping objective (25X NA 0.95) (bottom). In the former, the layer of Kwik-Sil glue is in the optical path. Scale bars 25 µm.

Next, to determine if we can image at high optical resolution over extended periods of time to achieve more sustained touch responses, we indented seedlings expressing the actin marker *pUBQ10::ABD2-mCherry*. As actin presents as a fine filamentous network, high-magnification imaging is needed to resolve fine features, and because actin dynamics are generally slower than calcium spikes, more prolonged imaging is required (42, 43, 48). In this experiment, we reversibly indent an epidermal cell in the hypocotyl with a 5 µm tungsten needle. This needle was used to more accurately mimic the experiments performed by Branco et al., where they showed that a smaller tip diameter resulted in more consistent actin reorganizations (42). On hypocotyl cells, the 5 µm tungsten needle can be used for reversible indentations due to a thicker epidermal cell wall (42). We observe that the actin cytoskeleton begins to accumulate at the indentation site (Figure 3b, Additional File 4). The response starts within 2 minutes, and a patch forms within 14 minutes. In accordance with the literature, we observe the actin patch formation only in 10-50% of the indentations per experiment (42, 43). It is currently unclear what causes this variability or stochasticity.

High-magnification, high-NA objectives, such as immersion objectives with a working distance of at least 150 µm, can be used with this device when it is placed on an inverted microscope. This ensures that we can resolve even subtle details at sub-cellular resolution, such as the actin filaments in this experiment. A larger working distance is required because the glue layer lies between the cover glass and the plant material. This also means that the glue layer is in the optical path and likely scatters light. Still, fine actin filaments can be resolved (Figure 3c). However, the signal that is collected can be significantly improved by using a water-dipping objective (Figure 3c). With a water-dipping objective, the objective is on the same side as the needle approaches the sample, and the glue layer is no longer in the optical path. Hence, there is less scattering and more signal collected. This especially improves imaging with marker lines with low expression or fine structures, as in this experiment.

### Applicability to other species and tissues

Thanks to the flexibility of the system, we can use it on different types of tissues and species. For example, the gametophyte of the fern *C. richardii*. This is a thin, quasi-two-dimensional structure because it mainly consists of a single cell layer. As such, like *A. thaliana* roots, this tissue can be easily approached from the side by the needle to puncture cells in different regions of the gametophyte. In the meristematic notch, where cell division occurs and cells are relatively young, puncturing a single cell with a 1 µm tungsten needle induces deformations throughout the surrounding tissue (Figure 4a, Additional File 5). We do not observe such extreme tissue deformations when puncturing a single cell in more mature tissue (Figure 4b, Additional File 6). These contrasting responses reflect differences in turgor pressure and mechanical properties across the tissue’s developmental stages, as previously described for this tissue by Woudenberg et al. (49).

**Figure 4.**
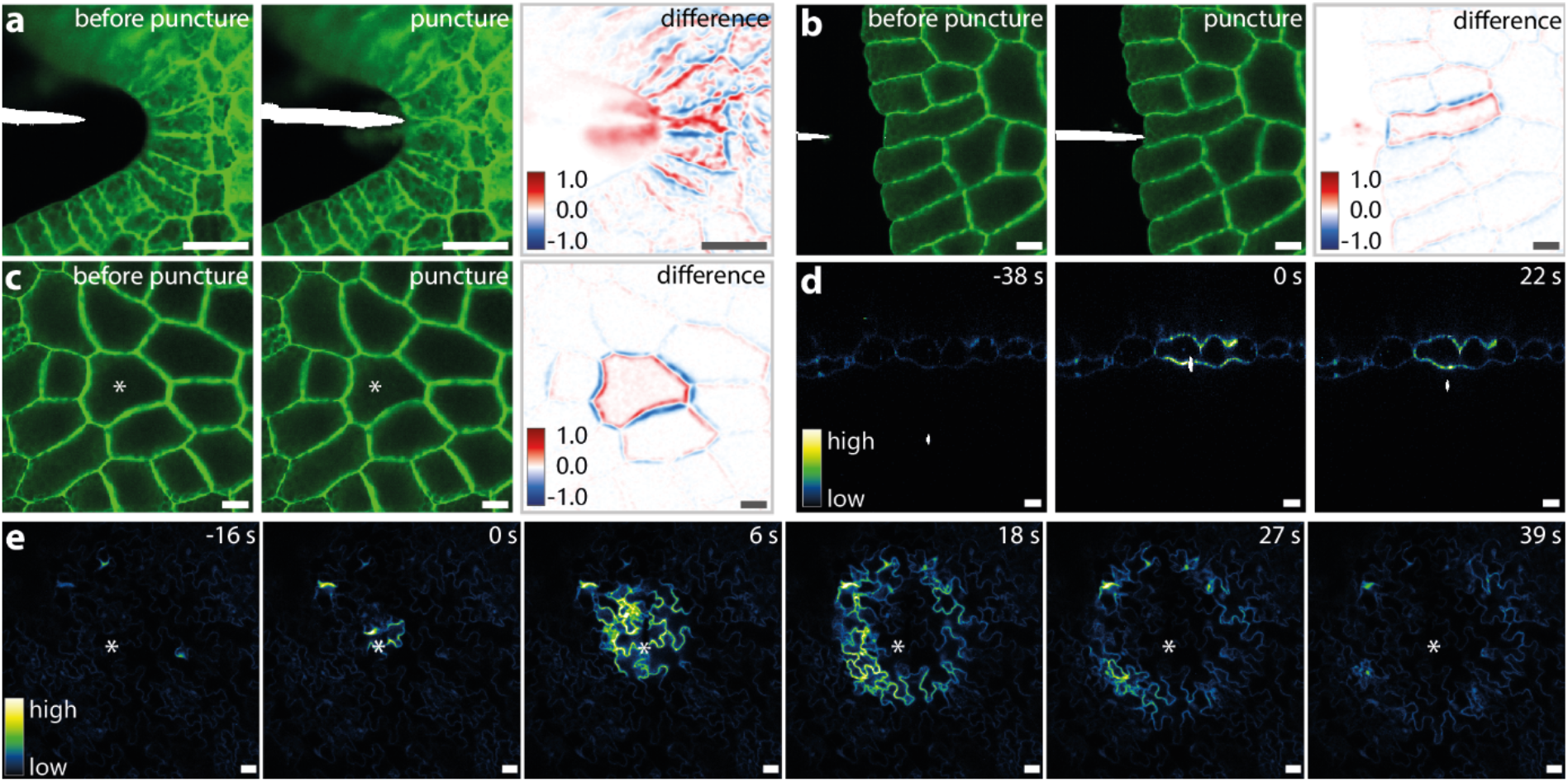
Indentation of different tissues. **(a-c)** Cells of *C. richardii* gametophyte stained with membrane stain LT-BDP (2) punctured with a 1 µm tungsten needle in the meristematic zone **(a)**, mature tissue on the side of the gametophyte **(b)** and mature tissue more centrally located **(c). (d-e)** Indentations of rosette leaf tissue of 3 week old *A. thaliana p35S::R-GECO1*.*2* plants. Water dipping objectives enable imaging of indentations of thicker or nontransparent tissues. Cross section imaging through xz-y imaging is useful to visualize indentation depth and cell deformations **(d)** while xy-z imaging can be used to study signaling perpendicular to the applied force **(e)**. The asterisks indicate the position of the tip of the needle in **c** and **e**. Scale bars: 25 µm

Since the tissue of the fern gametophyte is thin, it is also possible to target cells that are more centrally located. This is done by moving the needle horizontally above the sample to align the tip of the needle with a cell of choice, then moving the needle down to puncture it. In this case, DM shows that the punctured cell deforms due to a release of turgor pressure, while the cells surrounding the punctured cell show minimal deformation (Figure 4c, Additional File 7).

When tissues are thicker than a few cell layers and indentations must be performed on top of the tissue rather than from the side, inverted microscopes are no longer an option. These tissues must be imaged from the same side as the needle approaches them. This is possible with a water dipping objective on an upright microscope. To test this, rosette leaves of 3-week-old *A. thaliana* plants expressing *p35S::R-GECO1*.*2* were fixed to a small petri dish filled with water and indented with a 5 µm tungsten needle. Cross-sectional imaging in xz-y mode is used to visualize indentation depth and cell deformation more clearly than when the sample is imaged from the top in conventional xy-z mode (Figure 4d, Additional File 8). When imaging in xy-z mode, propagation of the calcium wave is observed over the whole tissue after the needle has touched the leaf (Figure 4e, Additional File 9). These results show that controlled indentations or tissue damage can also be performed on thicker, less transparent tissues.

### Functional imaging

The ability to perform high-resolution imaging with our indentation device allows it to be combined with more advanced imaging modalities, such as functional imaging. In this case, microscopy is used to make quantitative measurements of a physico-chemical property of interest, rather than solely imaging the localization of proteins or cellular components (50). To demonstrate the cross-compatibility of our device with functional Fluorescence Lifetime Imaging (FLIM), we indented seedling roots stained with the cell wall porosity probe CarboTag-BDP (1). This is a cell wall-binding fluorophore whose fluorescence lifetime changes in response to the nanoscale porosity of the cell wall (1). We perform reversible indentations with a 9 µm glass needle on epidermal cells in the maturation zone of *A. thaliana* seedling roots. As we indent the cell wall, we compress the carbohydrate meshwork. We expect this to deform like a sponge, expelling water and compressing the network, thereby leading to predictable changes in the fluorescence lifetime. As the network tightens and porosity decreases, the probe’s fluorescence lifetime increases. Indeed, we find that the fluorescence lifetime of the indented area is 25% higher than that at the same site before indentation and 25% higher than that of cell walls adjacent to the indentation site (Figure 5). This demonstrates how our modular device is compatible even with sophisticated high-resolution microscopy methods.

**Figure 5.**
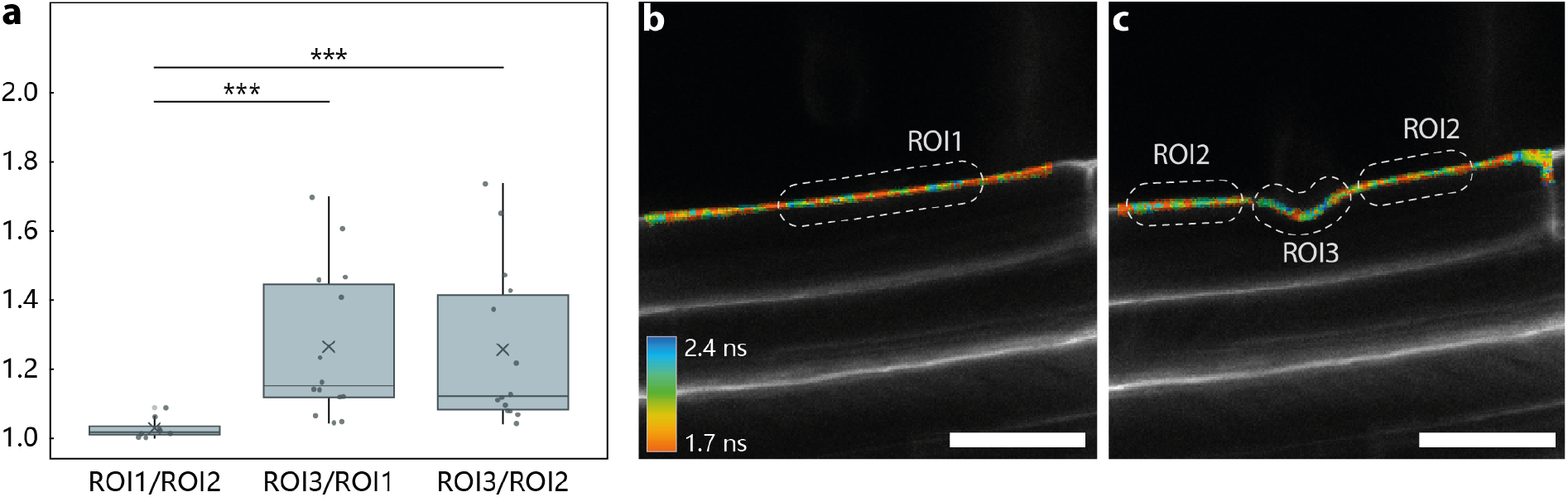
Cell wall properties during indentation can be visualized with functional imaging. **(a)** Cells in the maturation zone of *A. thaliana* seedlings stained with cell wall porosity probe CT-BDP (1) were indented with a 9 µm glass needle. **(a)** Quantification of fluorescent lifetime (τ) of CT-BDP in the cell wall, normalized to ROI1 (lifetime of the probe in the cell wall before indentation). Ratios of τ before indentation vs τ next to the indentation site (ROI1/ROI2), τ in the indentation site vs τ before indentation (ROI3/ROI1) and τ in the indentation site vs τ next to the indentation site (ROI3/ROI2) are depicted. n = 14, p < 0.001. **(b)** Image of the epidermal cell wall before indentation, where ROI1 indicates the region that was used for quantification of the fluorescent lifetime before indentation. Colors show fluorescent lifetime between 1.7 and 2.4 ns. **(c)** Image of the epidermal cell wall during indentation where ROI3 shows the region used for quantification of lifetime of the indentation site and the average of both ROI2 regions was used for quantification of the lifetime of the cell wall next to the indentation site.

## Discussion

In this study, we designed an indentation device that is low-cost and simple to build from primarily 3D-printed elements and some commercially available components. Due to the device’s modular design, it can be easily tailored to specific experimental requirements and different microscope setups. This flexibility makes it suitable for a wide range of applications, several of which are demonstrated in this paper.

A key advantage of the device is that cellular responses to mechanical stimulation can be imaged at high spatial and temporal resolution. This is not the case for all indentation devices or mechanobiology studies, where first a force is applied, and then the sample is transferred to a microscope (35). Important details of dynamic processes, or even whole rapid responses, are lost in this transfer period. Additionally, our use of piezo motors provides precise control over needle movement. This enables precise manipulation of single cells while imaging at high resolution.

We further demonstrate that needle geometry affects tissue manipulation: sharp needles with a tip diameter of 1 µm can be used to puncture single cells in the roots, whereas needles with a larger tip diameter can be used for reversible indentations. Our comparison of calcium responses in the root using a sharp needle versus a larger needle highlights the advantage of being able to exchange needles during experiments. This flexibility allows the study of cellular processes under different types of mechanical perturbation with the same setup, something that is not possible with laser ablation studies (21, 29), compression (29, 33-35), or stretching assays (36-38). The exchange of needles during the experiment revealed that some responses are amplified when the cell is damaged by puncturing versus milder indentations, suggesting that there may be synergistic effects between purely mechanical and DAMP-associated damage signaling.

We have demonstrated that a variety of tissues can be mounted and studied on this device. Thin or relatively transparent tissues can be easily studied with both upright and inverted microscopes. When indentations are applied to the epidermis from the side, as shown in the examples above, the tissue thickness is generally not a constraint, provided the needle can be moved as close to the cover glass as possible. However, for thicker, less transparent tissues that need to be indented from the top, an upright microscope equipped with a water dipping objective is necessary. When such an objective is available, both top and side indentations are possible. Moreover, the indentation site can be imaged without additional material in the optical path, reducing light scattering and potentially improving image quality. Being able to image leaf tissue is an advantage, since *N. benthamiana* leaf transiently expressing fluorescent proteins of interest is a widely used technique for studying protein localization (51, 52). These samples can be easily mounted and studied in our setup.

Compatibility with inverted microscopes is a significant practical advantage, as these systems are widely used. As shown here, the achievable resolution is sufficient to resolve fine structures such as thin actin filaments. Nevertheless, several limitations arise due to the glue used to fix the seedling. First, the glue layer is in the optical path when the device is used on inverted microscopes. The layer can scatter light, and the glue’s refractive index is unknown, potentially introducing refractive-index mismatches that reduce image quality. Optimal signal is obtained when imaged as close as possible to the glue layer. This requires some practice but is feasible. Second, oil- and water-immersion objectives with high NA typically have a working distance of 150 – 200 µm. While this is often sufficient to image through the glue layer into the sample, it limits the imaging depth and thereby constrains the range of possible indentation positions.

Another minor limitation is needle availability. In this paper, glass needles that have been manually pulled and blunted were used, which requires specialized equipment. Commercially available alternatives for needles are limited, yet acupuncture needles could also be used to perform reversible indentations and appear to be the best compromise if needle pulling equipment is not available. Finally, while the setup is compatible with most microscopes, it may not fit on all due to obstructions on the microscope body or differences in insert dimensions. However, the base designs can be adjusted with relative ease to fit those microscopes as well.

In conclusion, we present a modular, easily customizable indentation device suitable for a wide range of applications. The system can be assembled from 3D-printed parts and a small number of commercially available components. We showed that the device can be used for localized puncturing and reversible indentation of single cells. It can be used to study rapid as well as slow dynamic responses in thin and thick tissues. Moreover, the system can be combined with functional microscopy, providing further insights into plant cell wall properties during mechanical stimulation. At present, the system has primarily been used for qualitative analyses. The user can apply a well-defined mechanical deformation. These deformations can be quantified using differential maps. Yet, the exact magnitude of the force applied to the cell remains unknown. Incorporation of a sensitive load cell could enable more quantitative measurements. Additionally, the straightforward design could enable the mounting of different types of needles, such as injection needles. Overall, the design of this indentation device and its many applications are expected to make indentation experiments more accessible, thereby facilitating research in plant mechanobiology.

## Methods

### Design of the setup

The entire device consisted of five 3D printed elements, a block with 3 piezo motors and a controller from Thorlabs. All 3D printed components were designed in AUTODESK Tinkercad and printed with a Bambu Lab X1-Carbon 3D printer. Printing files were prepared in Bambu Studio with the following printing parameters: 0.2 mm layer thickness, 6 walls, 25% infill density and ironing of top surfaces. Black polylactic acid (PLA, Bambu Lab) filament was used as material for all elements of the setup, white breakaway filament for PLA (Bambu Lab) was used for the tree support structures of overhangs.

Part 1 of the 3D printed elements was a base that was adapted to specific microscope stages for Nikon (Additional File 10), TCS SP8 DIVE Leica (Additional File 11) and TCS SP8 or TCS SP5 Leica (Additional File 12). Part 2 was a circular sample holder that fit in the base. The cover glass or petri dish containing the sample could be placed into this sample holder after sample preparation. The sample holder with prepared sample could then easily be placed into base on the microscope when the whole setup is mounted. Four versions of the sample holder were designed: one where the complete cover glass is supported by the sample holder (Additional File 13), one that leaves more space for wide objectives (Additional File 14), one that lowers the sample closer to the objectives in case the working range of the microscope is not large enough (Additional File 15), and finally one that fits a 60 × 15 mm diameter petri dish (Greiner) instead of a cover glass for water dipping objectives (Additional File 16). Part 3, 4 and 5 made up the needle holder that could be mounted against the vertical piezo motor. Thin bars (Additional File 17) with a slit were designed to carry the needle. Any needle could be glued onto these bars with epoxy glue (RS Pro 132-605). These bars were clicked into the rotating part of the needle holder (Additional File 18) before or during the experiment. The angle of the needle could easily be adjusted due to the rotating property of this holder. The rotating holder was mounted onto the piezo motor using a 3D printed mount (Additional File 19) with M2 × 6 mm screws (RS Pro 293-280). An additional 6th element (Additional File 20) was designed to store the needles in 120.5 × 15.8 mm (width x height) square petri dishes (Corning). This element was glued into a petri dish with epoxy glue (RS Pro 132-605). Glued needles could be placed into the slits of the needle storage. Soft rubber was glued onto the lid of the petri dish to keep the needles in place.

For the block containing the motors of the setup, one piezo motor (Thorlabs, PD1/M – ORIC 20 mm linear stage with piezoelectric inertia drive – metric) was mounted vertically using a right-angle bracket (Thorlabs, PD1Z/M right-angle bracket adapter for 20 mm piezo stages – metric) with M4 × 6 mm (Thorlabs HW-KIT1/M) and M1.6 × 4 mm screws (provided with piezo motor) onto the XY piezo motor block (Thorlabs, PD1D/M – ORIC 20 mm monolithic XY stage with piezoelectric inertia drive – metric). This motor block was mounted into the slots of the 3D printed base for the microscopes with M4 × 10 mm screws (Thorlabs HW-KIT1/M). All three piezo motors were then connected to a KIM101 controller (Thorlabs, four-channel K-cube piezo inertia motor controller, with KPS201 power supply). The KIM101 controller could be connected through USB to a computer with the Thorlabs Advanced Positioning Technology (APT) User software version 3.21.6 for precise control of the needle.

### Needles

Seirin J-type no. 8 0.3 × 50 mm acupuncture needles, 5 µm and 1 µm tungsten needles (WPI, MN0010B and MN0010S respectively), and glass needles were glued onto separate 3D printed bars using epoxy glue (RS Pro 132-605). The glass needles were prepared from Drummond microcaps 50 µl glass capillaries and were pulled into needles using a Narishige PC-10 pipette puller with the following settings: heater level 60, weights 4, steps 1. The sharp tip of the needles was then melted using a microforge (Alcatel) to create round blunt needle tips of different sizes. To determine the tip diameter of the needles used in this study, brightfield images of the needles were acquired with a Nikon Eclipse Ti2 microscope (10X NA 0.25 air objective with 1.5x additional zoom). The circular selection tool in ImageJ version 1.52i was used to measure the diameter of the tip of the needles.

### Plant material and growth conditions

#### Arabidopsis thaliana

Seeds of Arabidopsis thaliana ecotype Columbia (Col-0), calcium reporter line p35S::R-GECO1.2 (47) and actin reporter line pUBQ10::ABD2-mCherry (53) were surface sterilized with a 70% ethanol solution for 3 minutes, rinsed with 96% ethanol solution and dried in sterile conditions for 1 hour. Sterile seeds were sown on ½ Murashige and Skoog (MS) agar plates and stratified at 4 °C for 2 or 3 days. The plates were then placed vertically in long-day light conditions (16h light – 8h dark) at 22 °C for 6 days. For rosette leaf material, seeds of p35S::R-GECO1.2 were sown and grown on soil in the greenhouse at 22 °C for 3 weeks.

#### Fern gametophyte

Sterilized Ceratopteris richardii spores (Hn-n strain) were grown on ½ MS agar plates with 1% sucrose in a Hettich MPC600 plant growth incubator at 28°C with 16h of 100 µmol m^-2^ s^-1^ as previously described (49). Gametophytes were grown from spores that were synchronized by imbibing the spores in the dark in water for at least 3 days. Hermaphroditic gametophytes were imaged after they developed a lateral notch meristem, approximately between 8-12 days after plating.

### Sample preparation for imaging

#### Arabidopsis thaliana seedlings

For functional imaging, wildtype seedlings were pre-stained in 10 µM CarboTag-BDP for 30 minutes (1). These seedlings were washed in ½ MS for 5 minutes before mounting. Other seedlings (wildtype, p35S::R-GECO1.2, pUBQ10::ABD2-mCherry) were used directly from the ½ MS plate. In all experiments, seedlings were fixed in place to be able to indent them without moving the seedling. To achieve this, a thin layer of Kwik-Sil low toxicity silicone adhesive (Kwik-Sil Adhesive World Precision Instruments Inc., Germany) was spread onto a 22 × 50 mm no. 1.5 cover glass, using leukopore tape (Duchefa) on the long sides of the cover glass as spacer to obtain an even glue layer. A seedling was placed on the glue 5 - 6 minutes after mixing the glue. For indentations of the hypocotyl cells, the cotyledons were cut off so that the hypocotyl could be placed onto the glue. Before placing the seedling, a medium reservoir was created with high vacuum grease (Dupont Molykote), into which ½ MS liquid medium or MilliQ with 10 µg/ml propidium iodide (Sigma-Aldrich) was pipetted after the seedling was placed onto the glue. The cover glass with the mounted sample was then placed into the 3D printed sample holder.

For actin imaging in the hypocotyl with the water dipping objective, a layer of Kwik-Sil adhesive was spread evenly using a cover glass onto the lid of a 60 × 15 mm round petri dish (Greiner). The seedling was placed onto the glue using tweezers after 3 minutes of mixing the glue. The petri dish was immediately filled with MilliQ and subsequently placed into the sample holder designed for petri dishes.

#### Fern gametophyte

Fern gametophytes were pre-stained overnight in ½ MS liquid medium containing 1 uM LipoTag-green (2). On the day of experiment, glue layer and reservoir were prepared on a coverglass as described above. Gametophytes were placed on the glue using P200 pipette tips and the reservoir was filled with ½ MS liquid medium.

#### Leaf tissue

Rosette leaves from 3 week old p35S::R-GECO1.2 plants were picked with tweezers. A layer of Kwik-Sil glue was spread evenly onto the lid of a 60 × 15 mm round petri dish (Greiner). The leaf was placed with the adaxial side in the glue layer. 5 minutes after mixing the glue, the petri dish was filled with MilliQ and placed into the sample holder that was designed for petri dishes.

### Imaging

#### Effect of different needles

Imaging was performed on a Nikon C2 inverted confocal laser scanning microscope. Propidium iodide was excited with a 561 nm laser at 7%, fluorescence was detected between 605 and 1000 nm. A 10X (NA 0.45) air objective was used with 1.5x zoom by using the intermediate magnification switch, and additional 2.5x zoom. Images of 512 × 512 pixels were acquired at 1 frame/sec. Timelapse imaging was performed at 1 fps while the needle was moved towards a cell, touched the cell for 5 seconds and was then retracted. After the needle was retracted from the cell, timelapse imaging continued for 2 minutes.

#### Calcium signaling in the root

Imaging was performed on a Nikon C2 inverted confocal laser scanning microscope. p35S::R-GECO1.2 was excited with a 561 nm laser at 8%, fluorescence was detected between 605 and 1000 nm. A 10X (NA 0.45) air objective was used with 1.5x zoom by using the intermediate magnification switch, and additional 3x zoom. Images of 512 × 512 pixels were acquired. Timelapse imaging was performed at 1 fps while the needle was moved towards a cell, touched the cell and was retracted from the cell again.

#### Actin reorganization

Imaging was performed on a Leica TCS SP8DIVE system. pUBQ10::ABD2-mCherry was excited with a 552 nm laser at 3.4%, fluorescence was detected between 570 and 641 nm. A 40X (NA 1.10) water immersion objective was used with 1.62x zoom. Images of 512 × 512 pixels were acquired. Z-stacks were made with 1 µm step sizes and timelapse imaging was performed continuously for 14 minutes. Actin imaging with the water dipping objective was performed on a Leica TCS SP5 system. pUBQ10::ABD2-mCherry was excited with a 561 nm laser at 10%, fluorescence was detected between 580 and 640 nm. A 25X (NA 0.95) water dipping objective was used with 3x zoom. Images of 512 × 512 pixels were acquired. Z-stacks were made with 1 µm step sizes and timelapse imaging was performed continuously for 15 minutes.

#### Fern gametophyte

Imaging was performed on a Nikon C2 inverted confocal laser scanning microscope. LipoTag-green (2) was excited with a 488 nm laser at 0.5%, fluorescence was detected between 500 and 550 nm. A 10X (NA 0.45) air objective was used with 2.6x zoom. Images of 512 × 512 pixels were acquired. Timelapse imaging was performed at 1 fps while the needle was moved towards a cell, punctured the cell and was moved away from the cell again.

#### Leaf indentation

Imaging was performed on a Leica TCS SP5 confocal laser scanning microscope. p35S::R-GECO1.2 was excited with a 561 nm laser at 6%, fluorescence was detected between 580 and 640 nm. A 25X (NA 0.95) water dipping objective was used with 1.5x (top view) or 1.7x (side view) zoom. Images of 512 × 512 pixels (top view) or 256 × 256 pixels (side view) were acquired. Timelapse imaging was performed with 1 fps (top view) or 0.5 fps (side view).

#### Functional imaging

Imaging was performed on a Leica TCS SP8 inverted confocal laser scanning microscope coupled to a Becker-Hickl SPC830 time-correlated single photon counting module. CarboTag-BDP (1) was excited with a 8% 488 nm laser line from a 40 MHz White Light pulsed laser, fluorescence was detected between 500 and 600 nm. A 63X (NA 1.20) water immersion objective was used with a 2.5x zoom. FLIM data for one 256 × 256 pixel image was acquired over 60 s at 400 Hz scan speed.

### Data analysis

ImageJ version 1.52i was used to process imaging data. For images where the needle is shown, the needle was segmented from the brightfield channels using ImageJ by applying a manual threshold.

#### Effect of the different needles

Images from the timelapses were saved as 8-bit individual tiffs without any adjustments to the images themselves. The following frames were then selected manually: 1 frame before touching the cell, 1 frame during touching the cell, 1 frame after the needle had been retracted. MATLAB 2023b was then used to generate difference maps of two comparisons: (i) before indentation versus during indentation, (ii) before indentation versus after retraction. Background signal was first removed from the selected frames by using a gaussian blur (σ1 Gaussian kernel) and subtraction of the mean background signal. Image registration was then performed to remove drift of the sample. Difference maps from the two selected frames were then generated by subtracting the two frames from each other and colored in blue/white/red. Script is supplemented (Additional File 21, Additional File 22). After analysis, images were cropped to 150 × 150 pixels. PI internalization and subsequent nuclear staining was monitored for each time-lapse using imageJ.

#### Calcium signaling

Brightness of the videos of the experiment with the 13 µm glass needle was increased by a factor 2.7. Such a brightness adjustment was not necessary for the videos of the experiments with the 1 µm tungsten needle.

#### Actin reorganization

Maximum projections were made from the z-stacks. Images were cropped to 380 × 380 pixels. For actin imaging with the water dipping objective, maximum projections were made from the z-stacks and one frame was selected for comparison to imaging on an inverted microscope.

#### Fern gametophyte

Images were cropped to 250 × 250 pixels. One image before indentation and one image after indentation was used for difference maps. Difference maps were generated as described above.

#### Leaf indentation

No adjustments in brightness were necessary. For the side view imaging, brightness was increased by a factor 7.3. The needle was segmented as described above.

#### Functional imaging

Acquired images were analyzed with SPCImage v.8.5 software to obtain two-component exponential decay curves for each pixel. After applying an intensity threshold, ROIs were selected. ROI1: before the indentation occurs, ROI2: the area next to the indentation site during indentation, ROI3: the indentation site. The ratios indentation site/before indentation (ROI3/ROI1) and indentation site/next to the indentation site (ROI3/ROI2) were used as a readout.

### Statistics

Pairwise Wilcoxon signed-rank test was used to calculate the significance between measurements of fluorescent lifetime of CT-BDP in the epidermal cell wall of the indented area versus the area next to the indentation and before indentation. These calculations were performed in R studio. P-values are mentioned in the caption of the figure.

## Declarations

### Ethics approval and consent to participate

Not applicable.

### Consent for publication

All authors approve publication of this manuscript.

### Availability of data and materials

The data associated with this manuscript will be made available upon request. The 3D design files (.stl) for the instrument, Matlab code for differential microscopy analysis and supplementary movies are publicly available at: https://git.wur.nl/bic/2026_plant_acupuncture_annadaamen

### Competing interests

We declare no competing interests.

### Funding

This work is funded by the European Research Council project Catch (project number 101000981) and the Dutch Research Council NWO through the Gravitation program GreenTE (project number: 024.006.001)

### Authors’ contributions

The idea for this work, its conceptualization and the design choices originated with AD, AB, CB and JS. AD designed needle mounts, sample holders and Leica SP5/SP8 insert. JS designed Nikon and Leica SP8 DIVE inserts. 3D printing was done by CH. AB manufactured glass needles. AD, CB and JS designed the experiments and contributed to data interpretation. AD and JL performed actin imaging. AD and SW performed experiment with *C. richardii*. All other experiments were performed by AD. MB analyzed FLIM data. JWB provided crucial help and feedback for experiments performed with the water-dipping objective. CB and JS supervised the project. AD, CB and JS wrote the manuscript. All authors reviewed and approved the final manuscript.

## Notes

https://git.wur.nl/bic/2026_plant_acupuncture_annadaamen

### Competing Interest Statement

The authors have declared no competing interest.

## References

1. Besten M, Hendriksz M, Michels L, Charrier B, Smakowska-Luzan E, Weijers D, et al. CarboTag: a modular approach for live and functional imaging of plant cell walls. Nat Methods. 2025;22(5):1081–90.

2. Besten M, Heesemans T, Froeling R, Zilliox M, Peeters Y, Romein R, et al. LipoTag: A minimal motif for live and functional imaging of plant cell membranes. bioRxiv. 2026:2026.04.08.717154.

3. Persat A, Nadell CD, Kim MK, Ingremeau F, Siryaporn A, Drescher K, et al. The mechanical world of bacteria. Cell. 2015;161(5):988–97.

4. Stones DH, Krachler AM. Against the tide: the role of bacterial adhesion in host colonization. Biochem Soc Trans. 2016;44(6):1571–80.

5. Tuson HH, Weibel DB. Bacteria-surface interactions. Soft Matter. 2013;9(18):4368–80.

6. Eloy C, Fournier M, Lacointe A, Moulia B. Wind loads and competition for light sculpt trees into self-similar structures. Nat Commun. 2017;8(1):1014.

7. de Langre E. Effects of Wind on Plants. Annual Review of Fluid Mechanics. 2008;40(Volume 40, 2008):141–68.

8. Argentati C, Morena F, Tortorella I, Bazzucchi M, Porcellati S, Emiliani C, et al. Insight into Mechanobiology: How Stem Cells Feel Mechanical Forces and Orchestrate Biological Functions. Int J Mol Sci. 2019;20(21).

9. Tassinari R, Olivi E, Cavallini C, Taglioli V, Zannini C, Marcuzzi M, et al. Mechanobiology: A landscape for reinterpreting stem cell heterogeneity and regenerative potential in diseased tissues. iScience. 2023;26(1):105875.

10. Anlas AA, Nelson CM. Tissue mechanics regulates form, function, and dysfunction. Curr Opin Cell Biol. 2018;54:98–105.

11. Hamant O, Haswell ES. Life behind the wall: sensing mechanical cues in plants. BMC Biol. 2017;15(1):59.

12. Gorelova V, Sprakel J, Weijers D. Plant cell polarity as the nexus of tissue mechanics and morphogenesis. Nat Plants. 2021;7(12):1548–59.

13. Tomobe H, Tsugawa S, Yoshida Y, Arita T, Tsai AY, Kubo M, et al. A mechanical theory of competition between plant root growth and soil pressure reveals a potential mechanism of root penetration. Sci Rep. 2023;13(1):7473.

14. Bronkhorst J, Kasteel M, van Veen S, Clough JM, Kots K, Buijs J, et al. A slicing mechanism facilitates host entry by plant-pathogenic Phytophthora. Nat Microbiol. 2021;6(8):1000–6.

15. Bronkhorst J, Kots K, de Jong D, Kasteel M, van Boxmeer T, Joemmanbaks T, et al. An actin mechanostat ensures hyphal tip sharpness in Phytophthora infestans to achieve host penetration. Sci Adv. 2022;8(23):eabo0875.

16. Ryder LS, Sprakel J, Talbot NJ. Mechanobiology of fungal invasion. Curr Biol. 2025;35(11):R485–R90.

17. Leger O, Garcia F, Khafif M, Carrere S, Leblanc-Fournier N, Duclos A, et al. Pathogen-derived mechanical cues potentiate the spatio-temporal implementation of plant defense. BMC Biol. 2022;20(1):292.

18. Zhou F, Emonet A, Denervaud Tendon V, Marhavy P, Wu D, Lahaye T, et al. Co-incidence of Damage and Microbial Patterns Controls Localized Immune Responses in Roots. Cell. 2020;180(3):440–53 e18.

19. Beauzamy L, Nakayama N, Boudaoud A. Flowers under pressure: ins and outs of turgor regulation in development. Ann Bot. 2014;114(7):1517–33.

20. Sampathkumar A, Yan A, Krupinski P, Meyerowitz EM. Physical forces regulate plant development and morphogenesis. Curr Biol. 2014;24(10):R475–83.

21. Louveaux M, Julien JD, Mirabet V, Boudaoud A, Hamant O. Cell division plane orientation based on tensile stress in Arabidopsis thaliana. Proc Natl Acad Sci U S A. 2016;113(30):E4294–303.

22. Hamant O, Heisler MG, Jonsson H, Krupinski P, Uyttewaal M, Bokov P, et al. Developmental patterning by mechanical signals in Arabidopsis. Science. 2008;322(5908):1650–5.

23. Mathew MM, Saccheri J, Das S, Rajagopalan K, Lane B, Ka S, et al. Wound repair in plants guided by cell geometry. Curr Biol. 2025;35(16):3851–68 e7.

24. Cameron C, Geitmann A. Cell mechanics of pollen tube growth. Curr Opin Genet Dev. 2018;51:11–7.

25. Reimann R, Kah D, Mark C, Dettmer J, Reimann TM, Gerum RC, et al. Durotropic Growth of Pollen Tubes. Plant Physiol. 2020;183(2):558–69.

26. Emonet A, Zhou F, Vacheron J, Heiman CM, Denervaud Tendon V, Ma KW, et al. Spatially Restricted Immune Responses Are Required for Maintaining Root Meristematic Activity upon Detection of Bacteria. Curr Biol. 2021;31(5):1012–28 e7.

27. Stockle D, Reyes-Hernandez BJ, Barro AV, Nenadic M, Winter Z, Marc-Martin S, et al. Microtubule-based perception of mechanical conflicts controls plant organ morphogenesis. Sci Adv. 2022;8(6):eabm4974.

28. Wang Z, Ye X, Huang L, Yuan Y. Modulation of morphogenesis and metabolism by plant cell biomechanics: from model plants to traditional herbs. Hortic Res. 2025;12(4):uhaf011.

29. Sampathkumar A, Krupinski P, Wightman R, Milani P, Berquand A, Boudaoud A, et al. Subcellular and supracellular mechanical stress prescribes cytoskeleton behavior in Arabidopsis cotyledon pavement cells. Elife. 2014;3:e01967.

30. Uyttewaal M, Burian A, Alim K, Landrein B, Borowska-Wykret D, Dedieu A, et al. Mechanical stress acts via katanin to amplify differences in growth rate between adjacent cells in Arabidopsis. Cell. 2012;149(2):439–51.

31. Basu D, Haswell ES. The Mechanosensitive Ion Channel MSL10 Potentiates Responses to Cell Swelling in Arabidopsis Seedlings. Curr Biol. 2020;30(14):2716–28 e6.

32. Mielke S, Zimmer M, Meena MK, Dreos R, Stellmach H, Hause B, et al. Jasmonate biosynthesis arising from altered cell walls is prompted by turgor-driven mechanical compression. Sci Adv. 2021;7(7).

33. Nakayama N, Smith RS, Mandel T, Robinson S, Kimura S, Boudaoud A, et al. Mechanical regulation of auxin-mediated growth. Curr Biol. 2012;22(16):1468–76.

34. Jacques E, Verbelen JP, Vissenberg K. Mechanical stress in Arabidopsis leaves orients microtubules in a ‘continuous’ supracellular pattern. BMC Plant Biol. 2013;13:163.

35. Louveaux M, Rochette S, Beauzamy L, Boudaoud A, Hamant O. The impact of mechanical compression on cortical microtubules in Arabidopsis: a quantitative pipeline. Plant J. 2016;88(2):328–42.

36. Bringmann M, Bergmann DC. Tissue-wide Mechanical Forces Influence the Polarity of Stomatal Stem Cells in Arabidopsis. Curr Biol. 2017;27(6):877–83.

37. Burian A, Hejnowicz Z. Strain rate does not affect cortical microtubule orientation in the isolated epidermis of sunflower hypocotyls. Plant Biol (Stuttg). 2010;12(3):459–68.

38. Robinson S, Huflejt M, Barbier de Reuille P, Braybrook SA, Schorderet M, Reinhardt D, et al. An Automated Confocal Micro-Extensometer Enables in Vivo Quantification of Mechanical Properties with Cellular Resolution. Plant Cell. 2017;29(12):2959–73.

39. Tanaka K, Choi J, Cao Y, Stacey G. Extracellular ATP acts as a damage-associated molecular pattern (DAMP) signal in plants. Front Plant Sci. 2014;5:446.

40. Tanaka K, Heil M. Damage-Associated Molecular Patterns (DAMPs) in Plant Innate Immunity: Applying the Danger Model and Evolutionary Perspectives. Annu Rev Phytopathol. 2021;59:53–75.

41. Zhang A, Matsuoka K, Kareem A, Robert M, Roszak P, Blob B, et al. Cell-wall damage activates DOF transcription factors to promote wound healing and tissue regeneration in Arabidopsis thaliana. Curr Biol. 2022;32(9):1883–94 e7.

42. Branco R, Pearsall EJ, Rundle CA, White RG, Bradby JE, Hardham AR. Quantifying the plant actin cytoskeleton response to applied pressure using nanoindentation. Protoplasma. 2017;254(2):1127–37.

43. Hardham AR, Takemoto D, White RG. Rapid and dynamic subcellular reorganization following mechanical stimulation of Arabidopsis epidermal cells mimics responses to fungal and oomycete attack. BMC Plant Biol. 2008;8:63.

44. Hoermayer L, Montesinos JC, Trozzi N, Spona L, Yoshida S, Marhava P, et al. Mechanical forces in plant tissue matrix orient cell divisions via microtubule stabilization. Dev Cell. 2024;59(10):1333–44 e4.

45. Zhang W, Kumar N, Helwig JR, Hoerter A, Iyer-Pascuzzi AS, Umulis DM, et al. Local traveling waves of cytosolic Ca(2+) elicited by defense signals or wounding are propagated by distinct mechanisms in Arabidopsis. Sci Signal. 2025;18(915):eadw2270.

46. Ma X, Hasan MS, Anjam MS, Mahmud S, Bhattacharyya S, Vothknecht UC, et al. Ca(2+) waves and ethylene/JA crosstalk orchestrate wound responses in Arabidopsis roots. EMBO Rep. 2025;26(12):3187–203.

47. Bellandi A, Papp D, Breakspear A, Joyce J, Johnston MG, de Keijzer J, et al. Diffusion and bulk flow of amino acids mediate calcium waves in plants. Sci Adv. 2022;8(42):eabo6693.

48. Qin L, Liu L, Tu J, Yang G, Wang S, Quilichini TD, et al. The ARP2/3 complex, acting cooperatively with Class I formins, modulates penetration resistance in Arabidopsis against powdery mildew invasion. Plant Cell. 2021;33(9):3151–75.

49. Woudenberg S, Plackett ARG, Hao Z, Suzuki H, Baez LA, Borassi C, et al. Transgenerational polarity axis inheritance during <em>Ceratopteris</em> embryogenesis. bioRxiv. 2025:2025.08.29.673061.

50. Besten M, Daamen A, Fendrych M, Borst JW, Sprakel J. Chemical Probes for Functional Plant Imaging. Annu Rev Plant Biol. 2026.

51. Goodin MM, Dietzgen RG, Schichnes D, Ruzin S, Jackson AO. pGD vectors: versatile tools for the expression of green and red fluorescent protein fusions in agroinfiltrated plant leaves. Plant J. 2002;31(3):375–83.

52. Guo Y, Bao Z, Deng Y, Li Y, Wang P. Protein subcellular localization and functional studies in horticultural research: problems, solutions, and new approaches. Hortic Res. 2023;10(2):uhac271.

53. Dyachok J, Sparks JA, Liao F, Wang YS, Blancaflor EB. Fluorescent protein-based reporters of the actin cytoskeleton in living plant cells: fluorophore variant, actin binding domain, and promoter considerations. Cytoskeleton (Hoboken). 2014;71(5):311–27.

