## Supplementary information for "Plant Acupuncture: a low-cost and open-source device for local mechanical stimulation"

Supplementary videos are available on:
https://git.wur.nl/bic/2026_plant_acupuncture_annadaamen

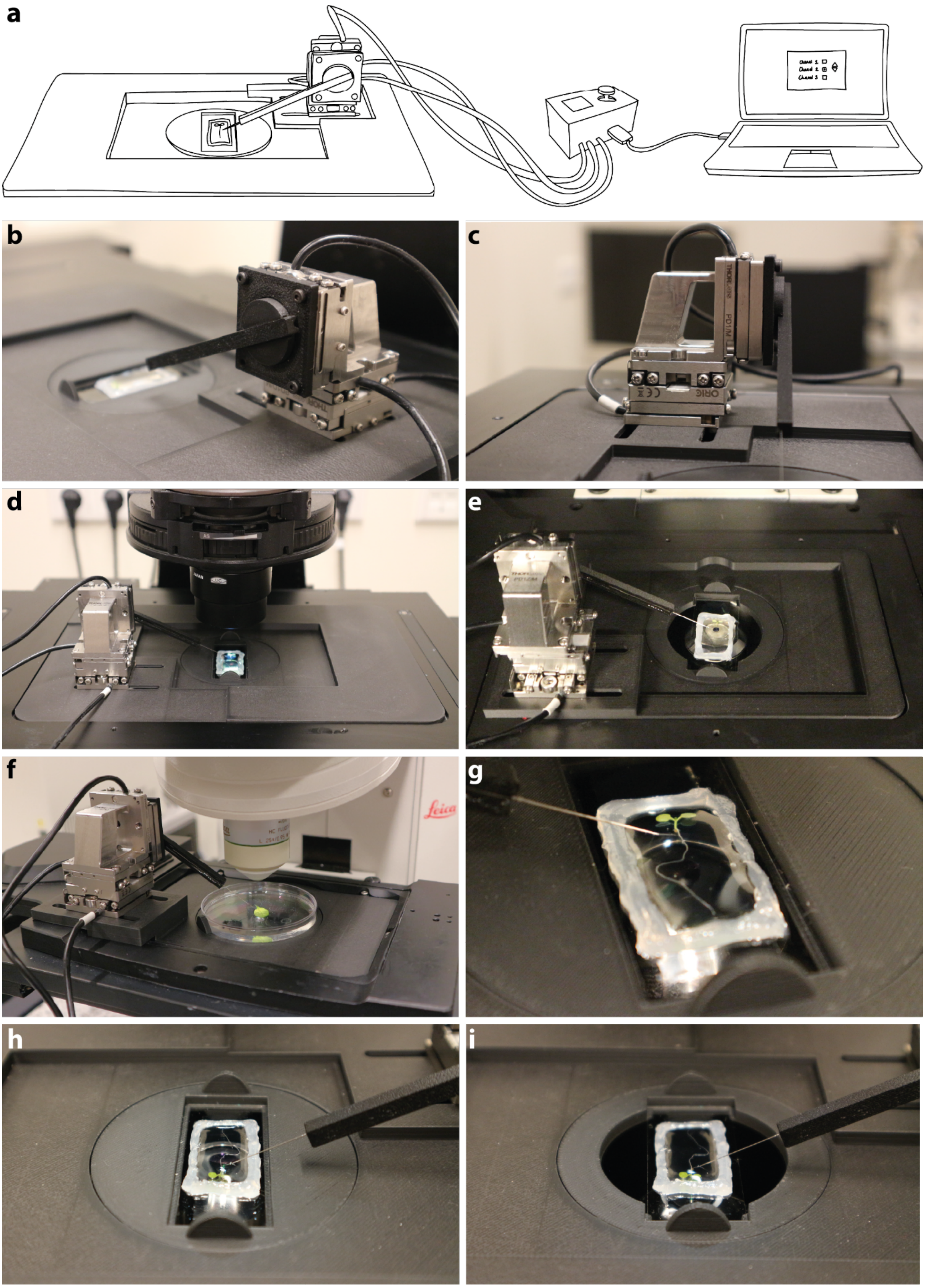
a
b
c
d
e
f
Leica
g
h
i

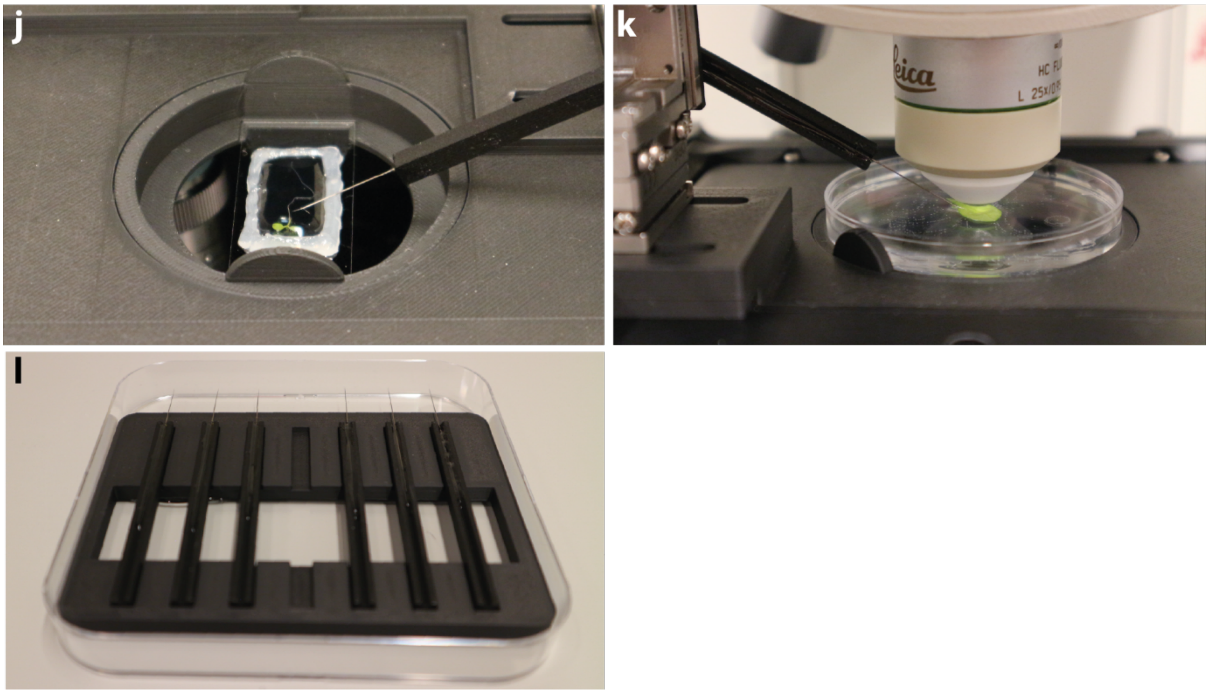


***Figure S1. Illustration and photos of the setup.*** **(a)** Schematic drawing of the complete setup with indentation device that is placed in the microscope, connected to the controller, which is connected to a laptop or PC with the software from the supplier. **(b-c)** Side views of the motor block that controls the needle. Consisting of a horizontal dual piezo motor for x-y movements and a vertically mounted piezo motor for z-movements. The needle is mounted with 3D printed elements onto the vertical piezo motor. **(d-f)** The setup in the Nikon **(d)**, SP8 DIVE **(e)**, and SP5 **(f)** microscopes. **(g)** Close-up of a sample prepared for inverted microscopes. **(h-k)** Close-up of sample holders designed for the bases, a stable holder that supports the entire cover glass **(h)**, a holder that leaves more space for larger immersion objectives **(i)**, a holder that lowers the sample more towards the objective for microscopes with a smaller working range **(j)**, a holder that fits a 6 cm petri dish for dipping objectives **(k)**. **(l)** 3D printed needle storage in a square plate.
